# Molecular basis of Cul3 ubiquitin ligase subversion by vaccinia virus protein A55

**DOI:** 10.1101/461806

**Authors:** Chen Gao, Mitchell A. Pallett, Tristan I. Croll, Geoffrey L. Smith, Stephen C. Graham

**Affiliations:** Department of Pathology, University of Cambridge, Tennis Court Road, Cambridge, CB2 1QP, United Kingdom; Cambridge Institute for Medical Research, University of Cambridge, Wellcome Trust/MRC Building, Cambridge CB2 0XY, United Kingdom

**Author notes:** To whom correspondence should be addressed: Stephen C. Graham: Department of Pathology, University of Cambridge, Tennis Court Road, Cambridge, CB2 1QP, United Kingdom.

**Keywords:** Poxvirus, innate immunity, viral immunology, E3 ubiquitin ligase, protein structure, structure-function, isothermal titration calorimetry, X-ray crystallography, BTB-Kelch

## Abstract

BTB-Kelch proteins are substrate-specific adaptors for cullin-3 (Cul3) RING-box based E3 ubiquitin ligases, which mediate protein ubiquitylation leading to proteasomal degradation. Vaccinia virus encodes three BTB-Kelch proteins, namely A55, C2 and F3. Viruses lacking A55 or C2 demonstrate altered cytopathic effect in cultured cells and altered pathology *in vivo*. Previous studies show that the ectromelia virus orthologue of A55, EVM150, interacts with Cul3 in cells. We show that A55 binds directly to Cul3 via its N-terminal BTB-BACK domain, and together they form a 2:2 complex in solution. The crystal structure of the A55/Cul3 complex was solved to 2.8 Å resolution. The overall conformation and binding interfaces resemble those of the cellular BTB-BACK/Cul3 complex structures, despite low sequence similarity of A55 to cellular BTB-BACK proteins. Surprisingly, despite this structural similarity the affinity of Cul3 for A55 is significantly higher than for reported cellular BTB-BACK proteins. Detailed analysis of the binding interface suggests that I48 from A55 at the BTB/Cul3 interface is important for this high-affinity interaction and mutation at this site reduced the affinity by several orders of magnitude. I48 is conserved only in close orthologues of A55 from poxviruses, but not in C2, F3, or other poxvirus or cellular BTB-Kelch proteins. The high affinity interaction between A55 and Cul3 suggests that, in addition to directing the Cul3-RING E3 ligase complex to degrade cellular/viral target proteins that are normally unaffected, A55 may also sequester Cul3 from cellular adaptor proteins and thus protect substrates of these cellular adaptors from ubiquitination and degradation.

---

Vaccinia virus (VACV)^1^ is a double-stranded DNA virus in the *Orthopoxvirus* genus of the *Poxviridae*. Historically VACV was used as the vaccine to eradicate smallpox (1). Its genome contains approximately 200 genes, about half of which are involved in the modulation of host immune response to viral infection, and the virus has been used as a model system to study innate immunity (2). The mechanisms by which several VACV proteins act to inhibit innate immune sensing and effector function, especially those involved in the inhibition of NF-κB signalling, have been well characterized (2,3). Nevertheless, many VACV immunomodulatory proteins are still poorly understood and one such protein is A55.

A55 is an intracellular protein encoded by the *A55R* gene of VACV (4). It belongs to the BTB (<u>B</u>ric-a-brac, <u>T</u>ramtrack and <u>B</u>road-complex)-Kelch protein family, which are substrate adaptor proteins specific for the cullin-3 (Cul3)-RING (Really Interesting New Gene) based E3 ubiquitin ligase (C3RL) complex (5). The N-terminal region of these proteins contains a BTB domain that mediates dimerization and binding to Cul3, a 3-box helical bundle region, and a BACK (for <u>B</u>TB <u>a</u>nd <u>C</u>-terminal <u>K</u>elch) domain that is likely responsible for correctly orientating the C terminus (5-13). The C-terminal region comprises 4-6 Kelch repeats arranged into a single β-propeller that captures the substrates for the C3RL complex; alternatively, these Kelch domains may also interact with actin filaments to regulate cytoskeleton organisation (5,9-11,14-19). In cells, there are many BTB-domain-containing proteins conjugated with different substrate recognition domains and their interactions with various substrates and C3RL complexes are implicated in several cellular processes including protein degradation, transcriptional regulation (Keap1), the gating of voltage-gated potassium channels (KCTDs), and cytoskeleton modulation (KLHLs) (19-25). Apart from mimiviruses, poxviruses are the only family of viruses that make BTB domain-containing proteins (26-30).

Deletion of A55 from VACV does not diminish virus replication in cultured cells (4). However, cells infected with VACV lacking A55 (vΔA55) demonstrated altered cytopathic effects, including the loss of Ca^2+^-independent cell adhesion and cellular projections, suggesting that A55 plays a role in the modulation of the cytoskeleton (4). The use of an intradermal murine model of infection demonstrated that infection with vΔA55 caused increased lesion size compared with wild-type virus, suggesting that A55 plays a role in altering the host immune response *in vivo* (4).

VACV encodes three BTB-Kelch proteins, namely A55, C2 and F3. Despite having similar domain organizations, A55 shares limited sequence similarity with C2 and F3 (22% and 25% aa identity, respectively). Like A55, C2 and F3 are dispensable for VACV replication in cultured cells (31,32). Infection of cells with vΔA55 or with VACV lacking C2 (vΔC2) produced a similar loss of Ca^2+^-independent cell adhesion, suggesting that A55 and C2 affect similar cellular pathways (4,31). However, intradermal infection *in vivo* with vΔC2 resulted in similar-sized lesions to wild-type infection but these lesions persisted for longer, distinct from the phenotype observed for vΔA55 (4,31). Infection with VACV lacking F3 (vΔF3) produced no distinct phenotype in cultured cells but intradermal infection yielded smaller lesions compared to wild-type virus (32). These results suggest that VACV BTB-Kelch proteins are functionally divergent despite having a conserved domain organization.

C3RLs are a family of multi-modular cullin-RING based E3 ubiquitin ligases that recruit substrates specifically via BTB-domain containing adaptor proteins (5,6). Cul3, the all-helical stalk-like scaffold subunit of C3RLs, interacts directly with BTB-domain-containing proteins via its N-terminal domain (6-8,13,24,33). The C-terminal domain of Cul3 interacts with the RING based E3 ligase protein to recruit the ubiquitin-loaded E2-conjugating enzyme for substrate ubiquitylation and is dispensable for binding to BTB-domain proteins (5,11,34). Crystal structures of several cellular BTB-domain proteins in complex with the Cul3 N-terminal domain (Cul3-NTD) have been reported (6,7,13,24). These structures revealed a unique mode of binding of BTB-containing adaptor proteins to the C3RL family of E3 ubiquitin ligases. Interaction with Cul3 is mainly via the BTB domain, with additional contacts from the 3-box region, while the BACK domain does not participate in the binding. The N-terminal 22 residues of Cul3 (N-terminal extension, or NTE) is usually disordered and dispensable for binding, and many reported binding studies of BTB-domain containing proteins to Cul3 were carried out with N-terminally truncated Cul3-NTD (Cul3_20-381_ for KLHL3, SPOP and KCTD5, Cul3_23-388_ for KLHL11 and Cul3_26-381_ for KEAP1)(6,7,13,24). However, the Cul3-NTE does provide extra hydrophobic contacts with the 3-box region upon binding to KLHL11 and KCTD5, resulting in significant increases in affinity (6,25).

Ubiquitin ligases act together with the proteasome to regulate the turnover of a large number of cellular proteins. Many viruses exploit the ubiquitylation-proteasomal degradation pathways to ensure successful infection and spread (35-41). To achieve this, viruses have evolved proteins that interact with ubiquitin ligase complex components to subvert the degradation pathways (35,37,39,42,43). The ectromelia virus (ECTV) orthologue of A55, EVM150, shares 93% sequence identity to A55. EVM150 has been reported to interact with Cul3 via its BTB domain and co-localizes with the C3RL and conjugated ubiquitin in cells (43). In addition, the BTB-domain of EVM150 was reported to inhibit NF-κB signalling, although Cul3 appeared dispensable for this activity (42).

In this study, we showed that A55 binds directly to Cul3 and solved the crystal structure of the complex. While the overall conformation of the A55/Cul3 complex is similar to reported cellular BTB/Cul3 structures, Cul3 binds A55 with much higher affinity than it does cellular BTB proteins. The non-conserved residue I48 at the BTB/Cul3 binding interface is required for this high affinity interaction. This strong A55/Cul3 interaction may allow VACV to redirect the E3 ubiquitin ligase complex to degrade novel target proteins and/or to subvert cellular BTB/Cul3 interactions to rescue proteins from degradation.

## Results

### A55 binds to Cul3 of the E3 ubiquitin ligase complex via its N-terminal BB domain

Poxvirus BTB-Kelch protein EVM150 has been shown to co-precipitate with Cul3 and modulate innate immune responses upon infection (42,43). A55 is also predicted to have a BTB-Kelch domain architecture and shares 93% aa identity to EVM150. To test whether A55 interacts with Cul3, co-immunoprecipitation experiments were performed using inducible HEK293T-REx cell lines expressing A55 with a FLAG-containing tandem affinity purification (TAP) tag at its N terminus (TAP-A55) or B14, an NF-κB inhibitor from VACV (44), with a C-terminal TAP tag (B14-TAP). Endogenous Cul3 co-immunoprecipitated with TAP-A55, but not B14-FLAG, when overexpressed in HEK293T cells (Fig. 1*A*). This suggests that Cul3 specifically co-immunoprecipitates with A55 and not with other VACV immune modulatory proteins. Furthermore, TAP-A55 co-immunoprecipitated with N-terminally myc-tagged Cul3 (myc-Cul3) but not with myc-Cul5, suggesting that A55 interacts specifically with Cul3 and not with other cullin family proteins (Fig. 1*B*). To dissect the region of A55 that binds to Cul3, the BTB-BACK (BB) and Kelch domains of A55 were tagged at the N terminus with TAP and were immunoprecipitated after overexpression in HEK293T-REx cells. Endogenous Cul3 co-immunoprecipitated with the N-terminal BB domain, but not with the C-terminal Kelch domain (Fig. 1*C*). These results suggest that, like the ECTV BTB-Kelch protein EVM150, A55 interacts with Cul3 and that this interaction is mediated solely by the N-terminal BB domain.

**Figure 1.**
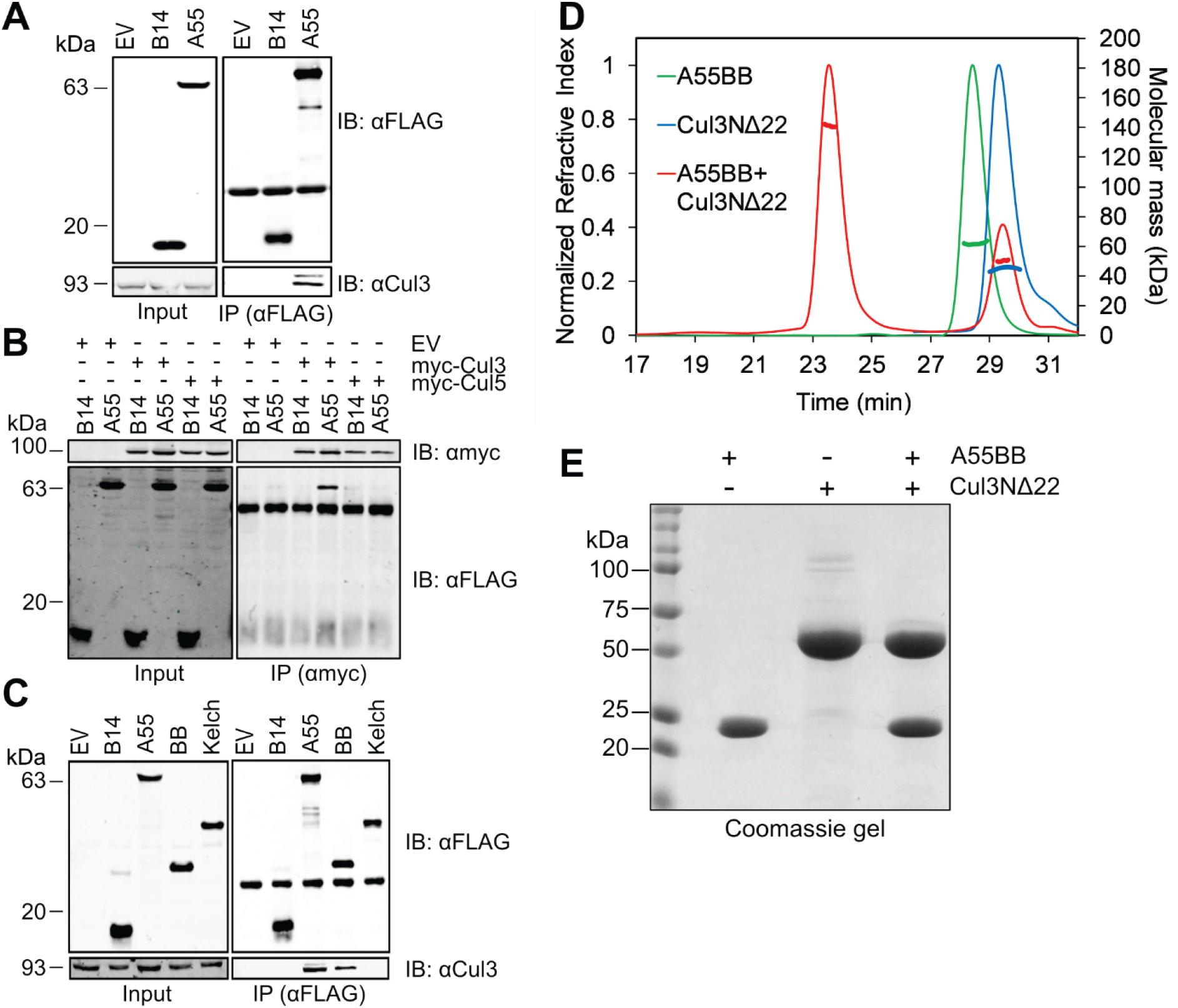
A55 binds to cullin-3 directly via its N-terminal BB domain. A-C. Representative immunoblots following immunoprecipitation (IP) of cleared lysates from HEK293T-Rex cell lines (**A**) expressing empty vector (EV), B14-TAP or TAP-A55, the TAP tag comprising STREP and FLAG epitopes; (**B**) expressing B14-TAP or TAP-A55 and transfected with plasmids encoding myc-Cul3 or myc-Cul5; (**C**) expressing EV, B14-TAP, TAP-A55, TAP-A55BB or TAP-A55 Kelch. Cells were lysed in (**A** and **C**) NP40 or (**B**) RIPA buffer. Immunoprecipitates were subjected to SDS-PAGE and immunoblotting. **A** and **C**. FLAG IP and immunoblotting for co-IP of endogenous Cul3. **B.** Myc IP and immunoblotting for co-IP of TAP-tagged B14 or A55. Input, cleared lysate. Data shown are representative of at least three independent experiments. **D.** SEC-MALS analyses showing the SEC elution profiles (thin lines) and molecular mass distribution (thick lines) across the elution peaks for A55BB (green, theoretical molecular mass 30 kDa, observed molecular mass 60 kDa), Cul3NΔ22 (blue, theoretical molecular mass 46 kDa, observed molecular mass 45 kDa) and A55BB and Cul3NΔ22 together (red, theoretic molecular mass 76 kDa, observed molecular mass 141 kDa) when eluting from a Superdex 200 10/300 GL column. **E.** Coomassie-stained SDS-polyacrylamide gel showing the species under each peak in (**D**).

### A55 is an obligate dimer in solution and forms a 2:2 complex with Cul3

Previous biochemical and structural analysis has shown that Cul3 binds cellular BTB-Kelch proteins via its N terminus (residues 1-388) while its C terminus (389-767) is not required for binding (6-8, 13). To test whether A55 forms a direct complex with Cul3 in solution, the A55 BB domain (A55BB, residues 1-250) and the Cul3 N-terminal domain (Cul3NΔ22, residues 23-388), were expressed in *E. coli* and purified according to protocols described in the experimental procedures. Size-exclusion chromatography coupled to multi-angle light scattering (SEC-MALS) studies together with SDS-PAGE analysis showed that A55BB exists as a homodimer in solution (expected molecular mass 60 kDa) (Fig. 1*D*). This is consistent with observations for other cellular BTB-BACK proteins (6-8). Cul3NΔ22 is monomeric (expected molecular mass 46 kDa). However, when A55BB and Cul3NΔ22 were mixed at approximately 1:1 molar ratio, a complex was formed with an apparent molecular mass of 141 kDa, consistent with a 2:2 complex of A55BB:Cul3N∆22 (expected molecular mass 152 kDa) (Fig. 1, *D* and *E*). Overall, the results show that A55 is dimeric in solution and binds directly to Cul3 to form a heterotetramer.

### A55 binds to Cul3 with low- to sub-nanomolar affinity

Isothermal titration calorimetry (ITC) experiments were carried out to determine the binding affinity between A55BB and Cul3. Two different truncations of Cul3 containing the N-terminal domain were used: Cul3N (residues 1-388) and Cul3NΔ22 (residues 23-388) to compare the binding affinities between A55 and Cul3 with or without the Cul3-NTE. A55BB formed 1:1 complexes with both Cul3NΔ22 and Cul3N with affinities in the low nanomolar (3 ± 1 nM) and sub-nanomolar (< 1 nM) range, respectively (Fig. 2, *A* and *B*; Table 1). The binding affinity of A55BB for Cul3N could not be determined accurately as rapid depletion of free Cul3N in the cell upon the introduction of A55BB prevented fitting of the resultant titration data to a single-site binding model. Attempts to lower the concentrations of A55BB and Cul3N or to use displacement titration experiments (45) were unsuccessful due to limitations of instrument sensitivity. Previous studies have shown cellular BTB-BACK proteins to bind the Cul3 N-terminal domain with much lower affinities than observed for A55BB (6-8, 13). To facilitate a direct comparison, the binding of KLHL3-BB to Cul3N and Cul3NΔ22 was measured by ITC (Fig. 2, *C* and *D*). These experiments confirmed that the affinity of Cul3 for A55 is approximately ten-fold tighter than for KLHL3 (Fig. 2, *C* and *D*; Table 1). Interestingly, the affinity of A55BB for Cul3NΔ22 was stronger despite the enthalpic contribution to the interaction (ΔH = -10.2 ± 1 kcal/mol) being lower than for the equivalent interaction between KLHL3-BB and Cul3NΔ22 (ΔH = -18.9 ± 1.3 kcal/mol) (Table 1). This suggests that the tighter interaction arises from a more favourable entropic contribution upon complex formation, such as the burial of exposed hydrophobic regions leading to the release of ordered solvent molecules and/or less conformational restriction of A55 upon complex formation. Taken together, the ITC data presented here show that the VACV A55 binds Cul3 more tightly than previously-studied cellular BB domain-containing proteins (Table 1) and that different thermodynamic properties of the interaction contribute to this enhanced binding.

**Figure 2.**
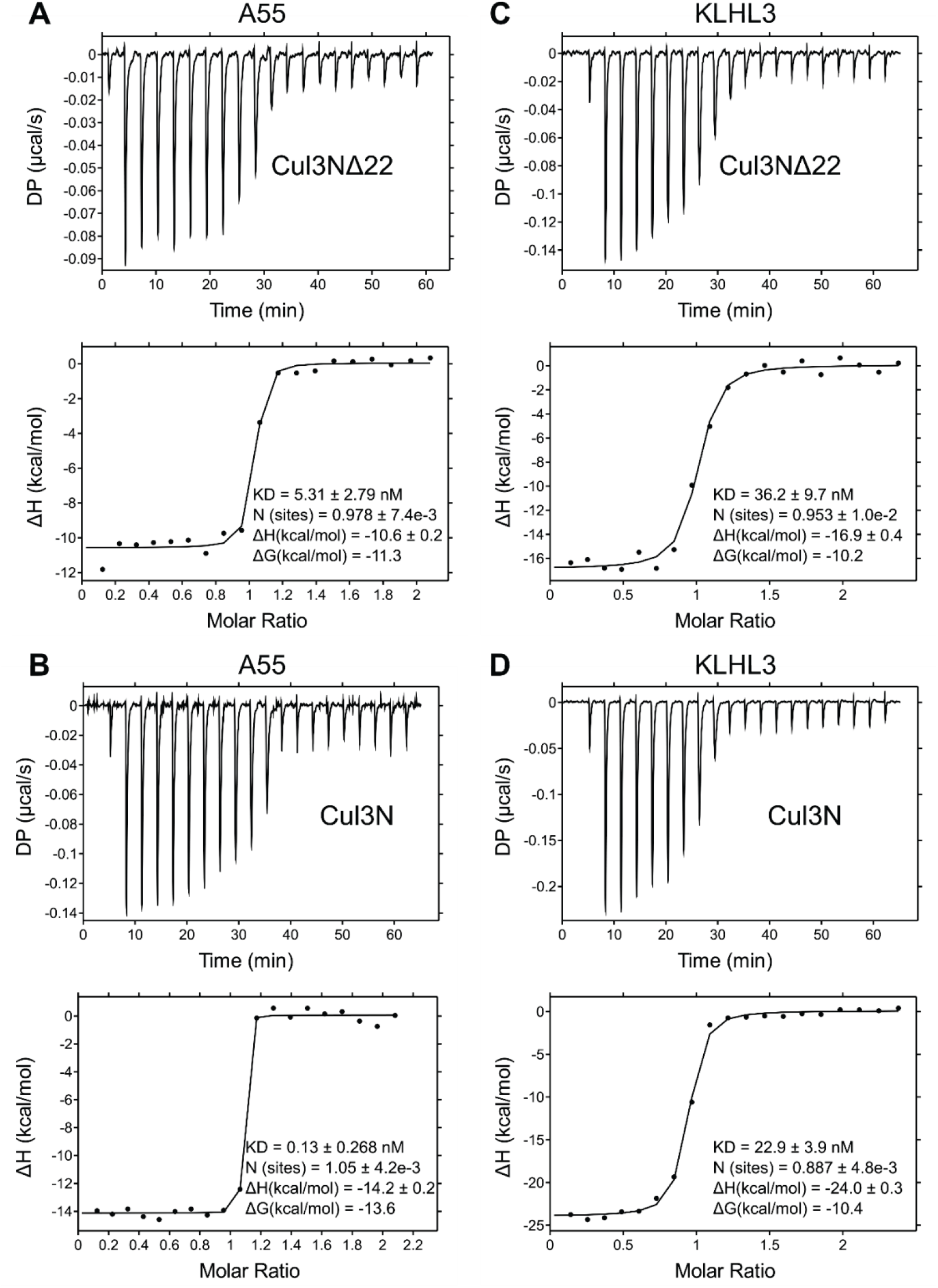
ITC studies show that A55 binds to Cul3 with nanomolar to sub-nanomolar affinity. A-D. Representative ITC titration curves showing interactions between A55BB and (**A**) Cul3NΔ22 or (**B**) Cul3N and between KLHL3, a human BTB related to A55, and (**C**) Cul3NΔ22 or (**D**) Cul3N. The top figure in each panel shows the baseline-corrected differential power (DP) versus time. The bottom figure of each panel is the normalized binding curve showing integrated changes in enthalpy (ΔH) against molar ratio. The corresponding dissociation constant (K_*D*_), number of binding sites (N), enthalpy change (ΔH) and change in Gibbs free energy (ΔG) for each representative experiment are shown. All experiments were performed at least twice independently.

**Table 1.**
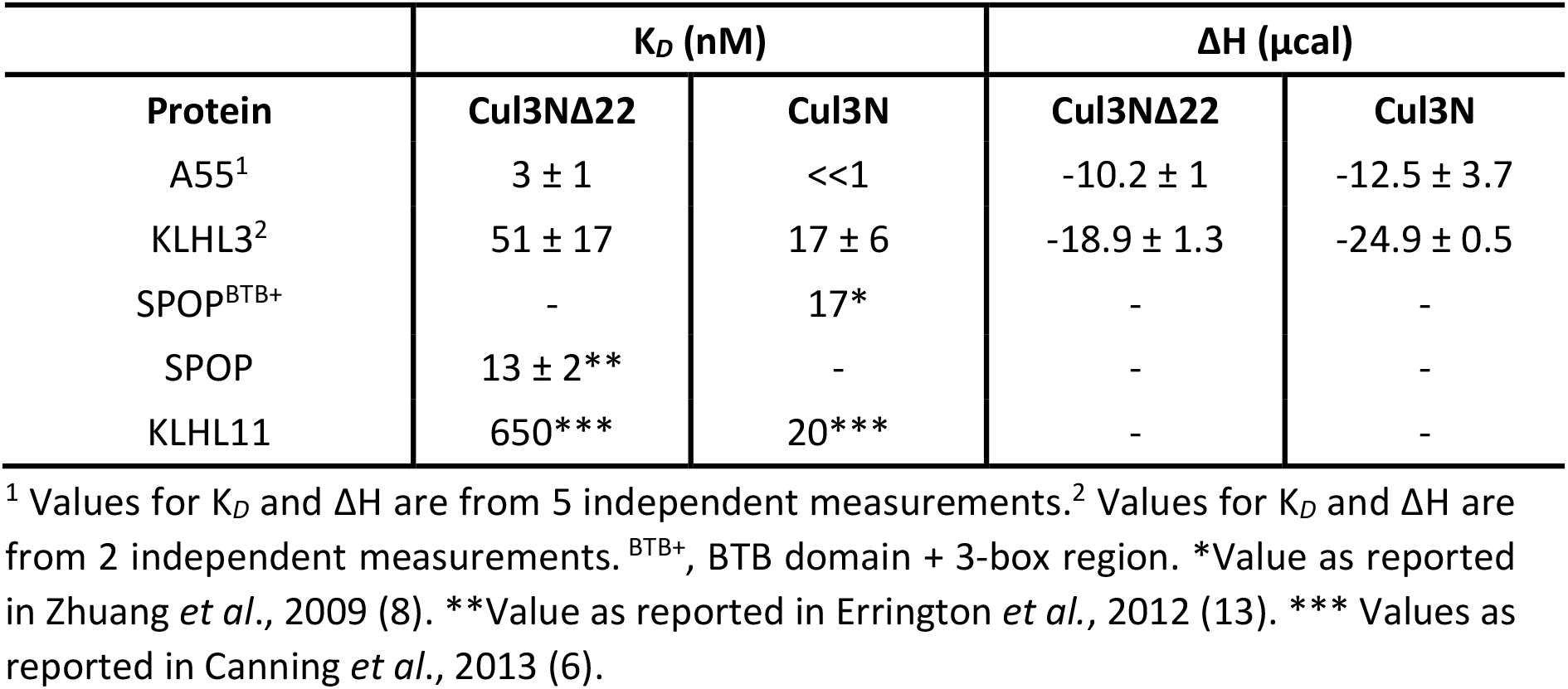
Comparison of the dissociation constants (K_*D*_) and enthalpic change (ΔH) for BB/Cul3 interaction. Experiments for this study were performed at least twice and mean ± SEM is shown. Experiments were performed *n* times and the mean ± SEM is shown.

| Protein | $K_D$ (nM) | | $\Delta H$ (μcal) | |
| --- | --- | --- | --- | --- |
| | Cul3NΔ22 | Cul3N | Cul3NΔ22 | Cul3N |
| A55<sup>1</sup> | 3 ± 1 | <<1 | -10.2 ± 1 | -12.5 ± 3.7 |
| KLHL3<sup>2</sup> | 51 ± 17 | 17 ± 6 | -18.9 ± 1.3 | -24.9 ± 0.5 |
| SPOP<sup>BTB+</sup> | - | 17* | - | - |
| SPOP | 13 ± 2** | - | - | - |
| KLHL11 | 650*** | 20*** | - | - |

<sup>1</sup> Values for $K_D$ and $\Delta H$ are from 5 independent measurements. <sup>2</sup> Values for $K_D$ and $\Delta H$ are from 2 independent measurements. <sup>BTB+</sup>, BTB domain + 3-box region. \*Value as reported in Zhuang *et al.*, 2009 (8). \*\*Value as reported in Errington *et al.*, 2012 (13). \*\*\* Values as reported in Canning *et al.*, 2013 (6).

### Determination of the A55BB/Cul3NΔ22 complex structure

To understand the molecular mechanism underlying the observed high-affinity interaction between A55 and Cul3, the A55BB/Cul3NΔ22 complex was purified and subjected to extensive crystallization screening for structural characterization. Initial trials did not yield any crystals. To promote crystallization, A55 was subjected to surface entropy reduction by reductive methylation (46) before being purified in complex with Cul3NΔ22 (Fig. S1). Crystals of methylated (M) A55BB in complex with Cul3NΔ22 grew as thin needles after two weeks. By using microseeding and varying the pH and concentration of the precipitants, optimized crystals were grown that diffracted to 2.3 Å in the best direction. Inspection of the diffraction data suggested severe anisotropy, with significantly worse diffraction along one axis (3.7 Å in direction 0.76 *a*^*^ - 0.65 *c*^*^) compared to the other major axes (2.6 Å in the direction b^*^ and 2.3 Å in the direction 0.92 *a*^*^ + 0.39 *c*^*^), so these data were processed with anisotropic scaling and truncation using STARANISO (47) and DIALS (48). The final processed dataset contained 23,509 unique reflections (Table 2), equivalent to the number of reflections expected for a 2.8 Å dataset collected from an isotropically-diffracting crystal with equivalent space group and unit cell dimensions. The anisotropy of diffraction was present in all crystals of the A55BB(M)/Cul3NΔ22 complex for which data were collected (>20 individual crystals).

**Table 2.** Data collection and refinement statistics.

| Data collection | A55BB Cul3NΔ22 complex |
| --- | --- |
| Beamline | Diamond I04 |
| Wavelength | 0.9795 Å |
| Resolution range (Å) | 116.6 - 2.3 (2.5 - 2.3) Å |
| Space group | C 1 2 1 |
| Unit cell (a/b/c) | 198.8/42.5/148.8 Å |
| Unit cell (β) | 128.4° |
| Total reflections | 300,453 (14,478) |
| Unique reflections | 23,509 (1172) |
| Multiplicity | 12.8 (12.4) |
| Completeness (spherical, %) | 53.2 (11.6) |
| Completeness (ellipsoidal, %) | 92.9 (86.4) |
| Mean I/sigma (I) | 15.3 (2.2) |
| R-merge | 0.087 (0.969) |
| CC1/2 | 1.000 (0.848) |
| Refinement |  |
| Resolution | 73.4 - 2.3 Å |
| Reflections used in refinement | 23,488 |
| Reflections in test (free) set | 1171 |
| R<sub>work</sub>/R<sub>free</sub> (%) | 28.0 / 29.6 |
| Protein residues | 539 |
| Number of non-hydrogen atoms | 4402 |
| RMS (bonds) | 0.012 Å |
| RMS (angles) | 1.5° |
| Ramachandran favoured* (%) | 96.1 |
| Ramachandran allowed* (%) | 3.6 |
| Ramachandran outliers* (%) | 0.4 |
| Rotamer outliers* (%) | 0.81 |
| Clashscore* | 1.6 |
| Average B-factor | 73.2 |

Statistics for the highest-resolution shell are shown in parentheses. R-values are as reported by autoBUSTER (51). \*Values as reported by Molprobit (65).

The structure of A55BB(M)/Cul3NΔ22 crystal was solved by molecular replacement using B-cell lymphoma 6 BTB domain (PDB code 1R29) (49) and Cul3_20-381_ from the SPOP/Cul3 complex structure (PDB code 4EOZ) (13) as the search models. While most of the Cul3NΔ22 molecule could be modelled with ease, the initial map for A55BB was less well-defined with relatively weak density for the 3-box and BACK regions. Anisotropic scaling of the diffraction data and the use of interactive molecular dynamics in ISOLDE (50) improved the model quality and fit to density significantly. The final model was refined using BUSTER (51) and has residuals R_work_/R_free_ of 0.280/0.296 with good overall geometry. Data collection and refinement statistics are summarized in Table 2. No crystals of A55BB(M) either alone or in complex with Cul3N could be obtained despite extensive crystallization trials.

### Structurally A55BB resembles cellular BTB-Kelch proteins with conserved Cul3-binding and dimerization interfaces

The structure of the A55BB(M)/Cul3NΔ22 complex contains one copy of each molecule in the crystallographic asymmetric unit (Fig. 3*A*). Consistent with the SEC-MALS analysis, a heterotetramer of A55BB(M)/Cul3NΔ22 can be observed by applying crystallographic 2-fold symmetry. A55 dimerization is mediated by the BTB domain, where the N-terminal helix (α1) forms a domain-swapped interaction with the symmetry-related molecule (Fig. 3*B*). The Cul3NΔ22 molecule is all-helical and closely resembles previously solved Cul3 N-terminal domain structures, with RMSD of 0.8 Å and 0.7 Å across 336 and 339 residues when aligned to the Cul3 structures in the KLHL3/Cul3NΔ19 (7) and KLHL11/Cul3NΔ22 (6) complexes, respectively (Fig. 3*C*). A55BB consists of a globular BTB domain (residues 1-118; helices α1-α6 and strands β2-β4) followed by a helix-turn-helix 3-box region (residues 119-149; helices α7-α8) and an all-helical BACK domain (residues 150-196, helices α9-α12) (Fig. 3*A*). A55BB closely resembles the equivalent regions of KLHL3 and KLHL11 (RMSDs of 2.2 Å and 2.5 Å across 167 and 181 residues, respectively), despite the low sequence conservation between A55 and these cellular proteins (Fig. S2 and Fig. 4), and the formation of dimers via an N-terminal helix domain swap is a conserved feature of all three proteins (Fig. 3*D*).

**Figure 3.**
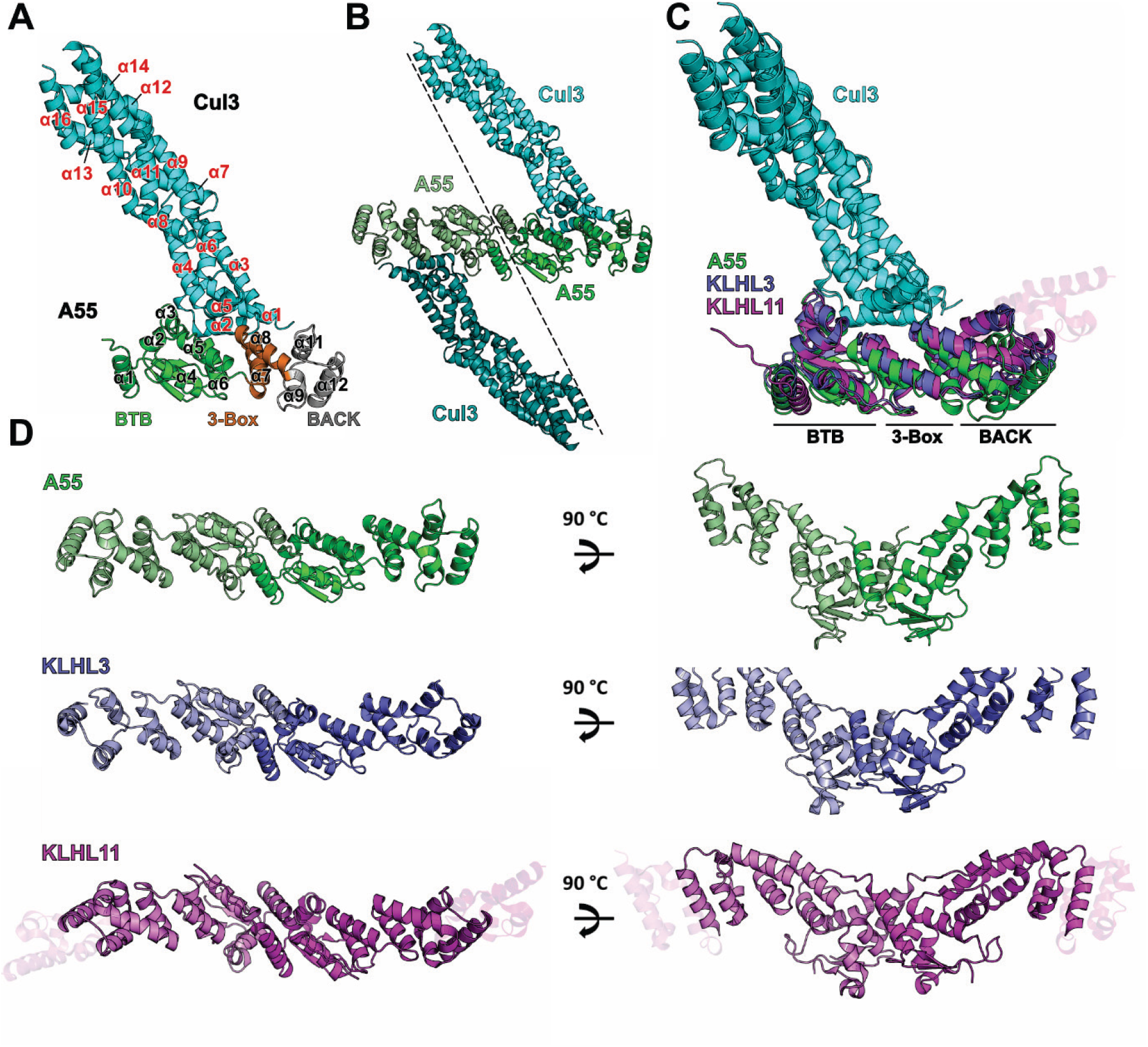
A55 and cellular BB domains share conserved modes of dimerization and Cul3 binding. **A.** Structure of the A55/Cul3NΔ22 heterodimer in the asymmetric unit as ribbon diagram. Cul3 is in cyan and the three domains of A55 (BTB, 3-box and BACK) are in green, orange and grey, respectively. Helices α1-α12 from A55 are labelled in black with the exception of α10, which is hidden behind α9 in the picture. Helices α1-α16 from Cul3 are labelled in red. **B.** A55/Cul3 dimer formed by applying crystallographic 2-fold symmetry. **C.** An overlay of three BTB-BACK/Cul3 complex structures (KLHL3/Cul3, PDB code 4HXI (7); KLHL11/Cul3, PDB code 4APF (6); A55/Cul3). The structures are aligned to the Cul3 part of the A55/Cul3 complex only. A55, KLHL3, KLHL11 and Cul3 are in green, purple, magenta and cyan, respectively, and the three sub-domains are marked. Additional helices at the C terminus of the KLHL11 BACK domain are shown as semi-transparent helices. **D.** Comparison of the dimers formed by A55, KLHL3 and KLHL11 BTB-BACK domains, colored as in (**C**).

**Figure 4.**
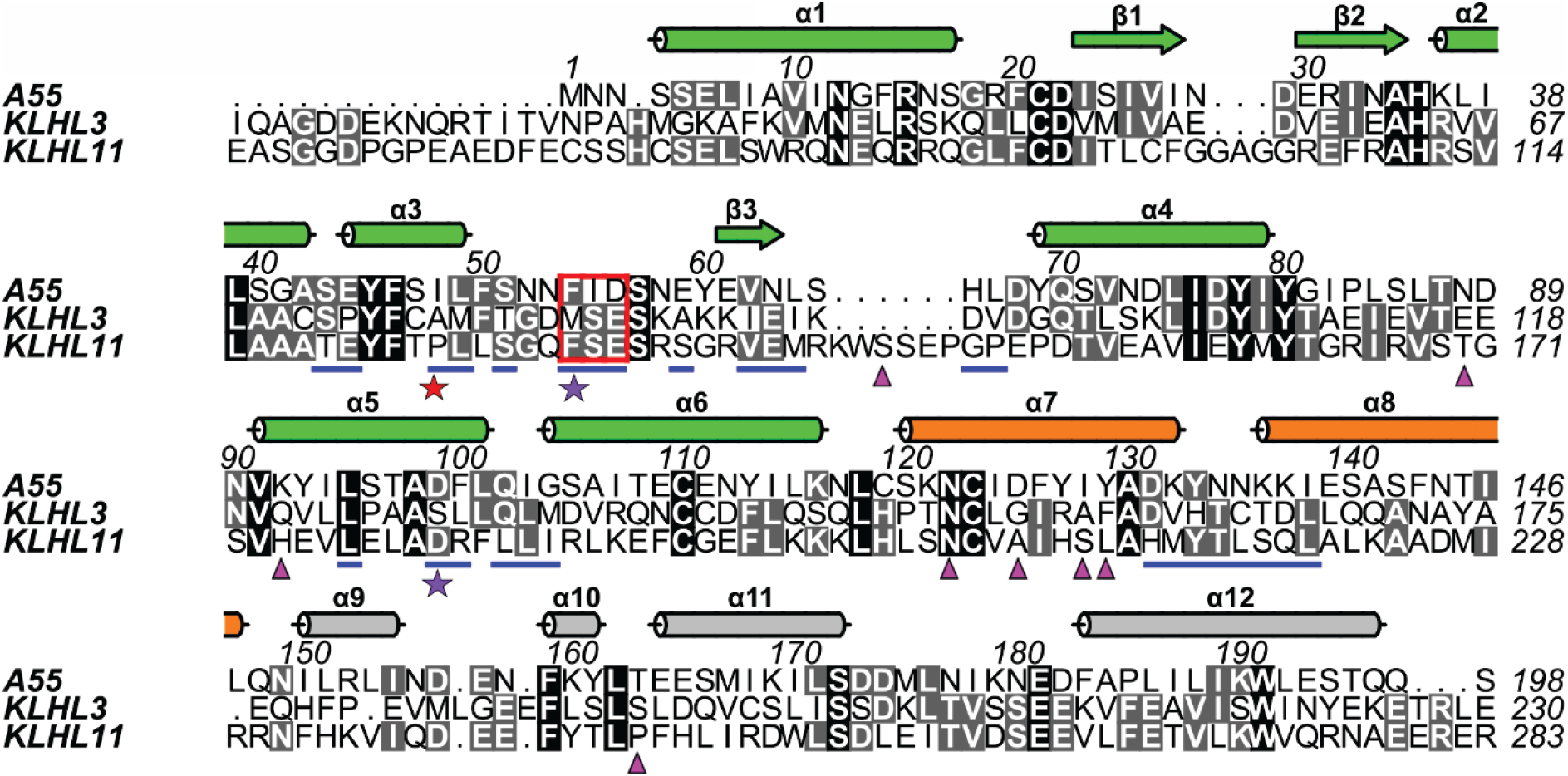
Structure-based sequence alignment of the A55, KLHL3 and KLHL11 BTB-BACK domains. Columns are shaded based on amino acid similarity. Secondary structural elements for A55 are shown above the aligned sequences and colored as in Fig. 3*A*. Residues at the A55/Cul3 interface are underlined in blue. Residues selected for subsequent mutagenesis studies of A55 are marked by stars at the bottom: the two conserved sites (F54 and D56) are marked by purple stars, while the non-conserved site (I48) is marked by a red star. Residues from KLHL11 that are involved in Cul3-NTE binding are marked by magenta triangles.

Overall, the A55BB/Cul3NΔ22 complex closely resembles other structures of Cul3 in complex with cellular BTB-domain containing proteins (Fig. 3*C*). The A55-binding interface of Cul3 is formed primarily of residues in helices α2 and α5, with extra contacts from the α1-α2 loop and from the C terminus of α3. Residues at the Cul3-binding interface of A55 are primarily found in the BTB domain with additional contacts in the 3-box region; the BACK domain does not contribute to the interaction (Fig. 4). This mode of interaction is consistent with the KLHL3/Cul3 (7), KLHL11/Cul3 (6) and KEAP1/Cul3 (PDB code 5NLB) complex structures (Fig. S2). Analyses of the interface areas and the number of interface residues for A55 and all available BTB/Cul3 complex structures showed no striking overall differences (Fig. S2). However, there appears to be a greater number of hydrogen bonds and less contribution from hydrophobic interaction in the A55BB/Cul3 complex when compared to the other BB/Cul3 structures (Fig. S2).

Only 196 of the 250 aa residues that comprise A55BB could be modelled confidently: the density for side chains in the last modelled helix of the BACK domain (α10, residues 180-196) is weak compared with density for side chains at the BTB/Cul3 interface (Fig. S3, *A* and *B*), and density for BACK domain residues 197-250 was not sufficiently well resolved to be modelled unambiguously (Fig. S3*C*). Correspondingly, the B factors of A55 residues at the Cul3-binding interface are lower than in the BACK domain (Fig. S3*D*). Inspection of the crystal lattice shows large solvent channels next to the BACK domain and this lack of crystal contacts at the C terminus is likely to account for the poor density observed in this region (Fig. S3, *C* and *E*). When superimposing KLHL11 and KLHL3 onto different regions of A55, inter-domain flexibility is evident (Fig. S4, *A* to *D* and *E* to *F*, respectively). Two pivot points in the BB structure can be found: the BTB_α5-α6_ helix-turn-helix and the 3-box region, respectively; the latter appears to be the major pivot point around which the BTB and BACK domain rotate relative to each other (Fig. S4, *I* to *K*). When measuring the angles between different subdomains (BTB_α1-4_, BTB_α5-α6_, 3-box and BACK), the angle formed by BTB_α5-α6_ -3-box-BACK in A55 is much larger compared to the corresponding angles in KLHL3 and KLHL11, thus rendering A55 more linear across the BB domain than KLHL3 and KLHL11 (Fig. S4, *I* to *K*).

Crystals of the A55BB/Cul3NΔ22 could be obtained only when the A55BB protein had been methylated *in vitro*. While there was density consistent with the presence of additional atoms adjacent to the amino termini of two lysine side chains (K36 and K132), they were not sufficiently well resolved to allow modelling of the methyl groups (Fig. S5*A*). Only one lysine (K136) was found at the binding interface (Fig. S5*B*) and ITC studies showed that methylated A55BB binds to Cul3 with affinity similar to unmodified A55BB (Fig. S5, *C* and *D*).

### The non-conserved residue I48 is required for the high-affinity binding of A55 to Cul3

Despite similarity in the overall structures, A55 binds Cul3 with much higher affinity than other BTB proteins. It has been suggested the key determinant for the interaction with Cul3 is a conserved φ-X-D/E motif found in the α3-β4 loops of the BTB domain, where φ is a hydrophobic residue and X is any residue (6,7,13). This motif exists in A55, corresponding to residues F54 (φ), I55 (X) and D56 (D/E), respectively (Fig. 4 and Fig. 5*B*). As in KLHL3, KLHL11 and SPOP, the side chain of residue φ (F54) in A55 is buried in a hydrophobic cavity on the surface of Cul3 (Fig. 5, *C* to *F*). Mutation of the ϕ residue in SPOP to a charged residue (M233E) completely abolished binding to Cul3, highlighting the significance of the φ residue for the interaction (13). An F54E mutant of A55BB was purified and shown to have the similar thermal stability to the wild-type protein (Fig. 6, *A* and *B*). ITC analysis demonstrated that the F54E mutation reduces the affinity of A55BB for Cul3NΔ22 and Cul3N by approximately 7 to 20 fold compared to the wild-type protein, yielding binding constants (K_*D*_s) similar to those of cellular BTB proteins (Fig. 6, *C* and *D*; Table 3). This suggests that φ residue F54 of A55 is involved in the interaction but not critical for binding to Cul3. A55 residue D56, equivalent to the D/E residue of the φ-X-D/E motif, forms side chain and backbone hydrogen bonds with Cul3 residues S53 and F54, respectively. A D56A mutation was introduced into A55 but the mutant could not be purified following bacterial expression, suggesting D56 is critical for the correct folding of A55.

**Table 3.**
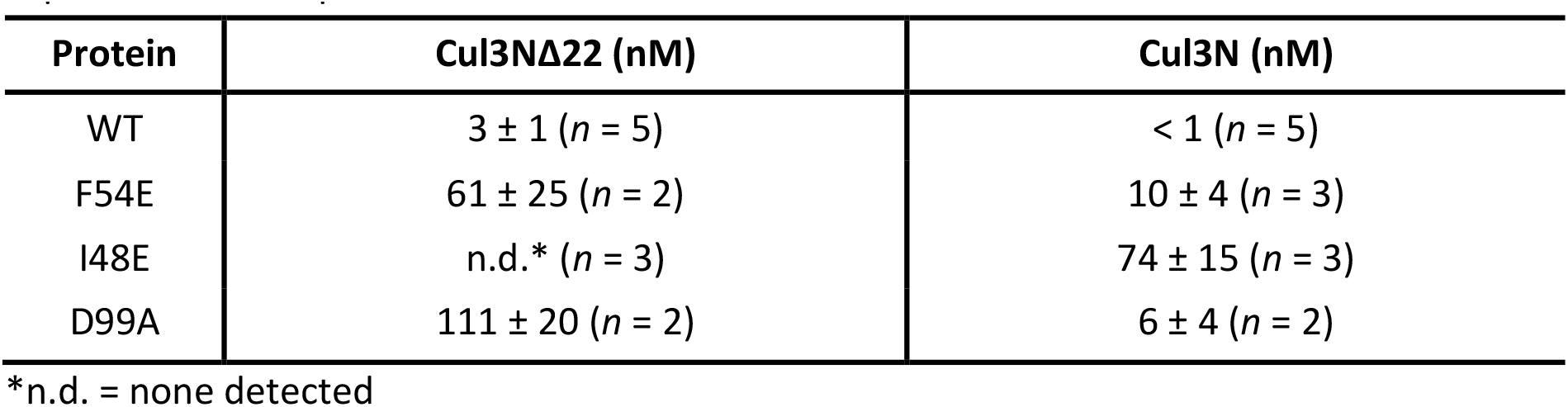
Comparison of the dissociation constants (K_*D*_) for WT and mutant A55BB for Cul3. Experiments were performed n times and the mean ± SEM is shown.

**Figure 5.**
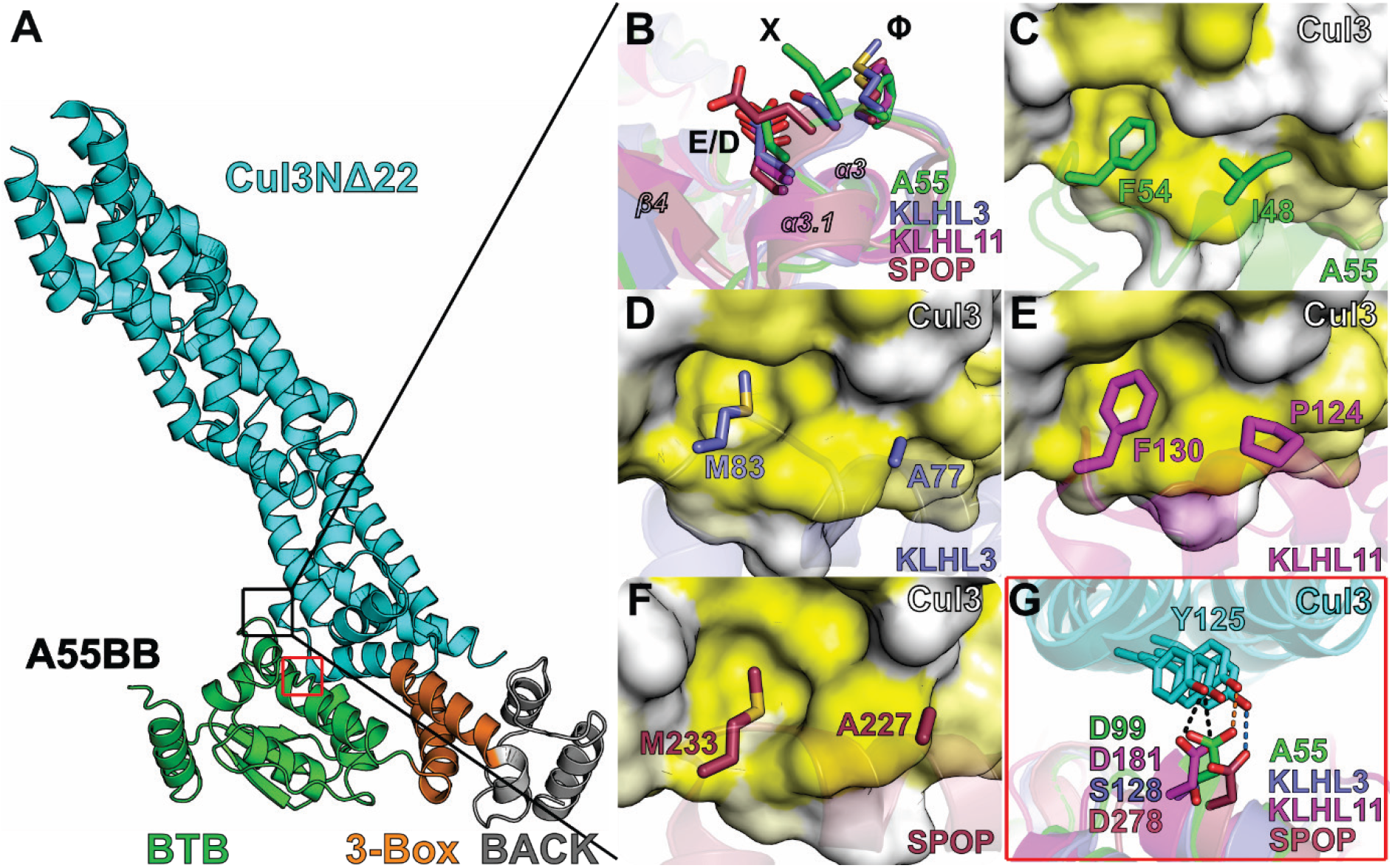
Conserved and non-conserved interactions at the interface between A55 and Cul3. **A.** The A55BB/Cul3NΔ22 complex structure with two key Cul3 binding sites in the BTB domain boxed in black (enlarged in **B** to **F**) and red (enlarged in **G**). **B.** Structural overlay of the φ-X-D/E motifs from A55, KLHL3, KLHL11 and SPOP. **C**-**F.** Surface of Cul3 colored by residue hydrophobicity from yellow (hydrophobic) to white (polar) (70). Hydrophobic binding pockets are shown for F54 of A55, M83 of KLHL3, F130 of KLHL11, and M233 of SPOP, which are equivalent to the φ residue of the φ-X-D/E motif, and for I48 of A55 and its equivalent residues A77, P124 and A227 in KLHL3, KLHL11 and SPOP, respectively. **G.** An overlay of the hydrogen bond formed between Y125 of Cul3 and D99 of A55 with equivalent residues S128, D181 and D278 in KLHL3, KLHL11 and SPOP, respectively.

**Figure 6.**
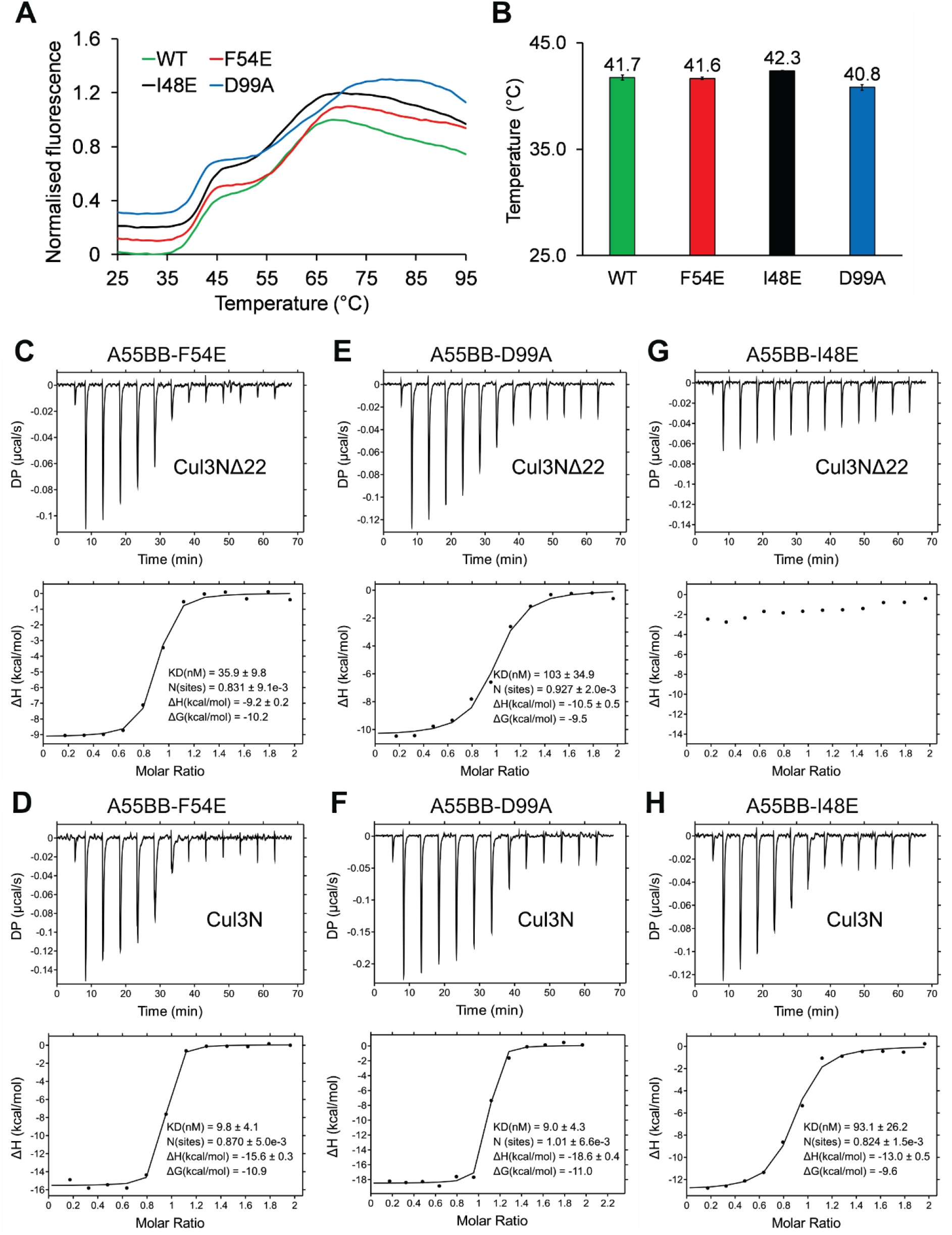
I48E mutation significantly impaired A55 binding to Cul3. **A.** Representative thermal melt curves of wild-type A55, F54E mutant, I48E mutant and D99A mutant from differential scanning fluorimetry (DSF) studies. Curves are offset along the vertical axis for clarity. All experiments were performed in triplicate. **B.** A comparison of the melting temperatures for wild-type A55 (green), F54E (red), I48E (black) and D99A (blue) mutants. Error bars show the standard errors of the mean from experiments performed in triplicate. **C-D.** Representative ITC titration curves showing A55BB mutant F54E binding to Cul3NΔ22 and Cul3N, respectively. **E-F.** Representative ITC titration curves showing A55BB mutant D99A binding to Cul3NΔ22 and Cul3N, respectively. **G-H.** Representative ITC titration curves showing A55BB mutant I48E binding to Cul3NΔ22 and Cul3N, respectively. The top figure in each panel shows baseline-corrected differential power (DP) versus time. The bottom figure of each panel is the normalized binding curve showing integrated changes in enthalpy (ΔH) against molar ratio. The corresponding dissociation constant (K_D_), number of binding sites (N), enthalpy change (ΔH) and change in Gibbs free energy (ΔG) for each representative experiment are shown. All experiments were performed at least twice independently.

As the φ-X-D/E motif was not absolutely required for binding of A55 to Cul3, the contribution of other residues was investigated. A hydrogen bond is formed between D99 of A55 and Y125 of Cul3 and this interaction is conserved in KLHL11 and SPOP but not in KLHL3 (Fig. 5*G*). Mutation at this site only caused moderate reduction in affinities for both Cul3NΔ22 and Cul3N (Fig. 6, *E* and *F*). Residue I48 of A55 is adjacent to the φ residue F54 and, like F54, the side chain of I48 extends deep into a hydrophobic cleft on the Cul3 surface (Fig. 5*C*). This residue is not conserved in KLHL3, KLHL11 or SPOP and the equivalent residues (A77, P124 and A227, respectively) have smaller side chains and form less extensive interactions (Fig. 5, *D* to *F*). An A55BB-I48E mutant was generated and shown to have similar thermal stability to the wild type protein (Fig. 6, *A* and B). Surprisingly, the I48E mutation completely abolished binding to Cul3NΔ22 and reduced the affinity for Cul3N by several orders of magnitude (Fig. 6, *G* and *H*) to well-below those of the cellular BTB proteins for Cul3 (Table 1 & 3). This suggests that residue I48, which is not conserved in cellular BTB proteins, forms a key hydrophobic interaction that mediates the high-affinity interaction between A55 and Cul3. The residual binding to Cul3N is likely to be mediated via contacts with Cul3N-NTE, as has been characterized previously for KLHL11 (6).

## Discussion

An interaction between a poxvirus BTB-Kelch protein and Cul3 has been demonstrated previously for the ECTV protein EVM150 by co-immunoprecipitation from transfected cells (42,43). Cellular BTB proteins have been reported to bind directly to Cul3 (6-8,13,24,25,33). However, due to low sequence identity (20–25%) between poxvirus and cellular BTB proteins (Fig. 4), it was unclear whether EVM150 and other poxvirus BTB proteins would bind Cul3 in a similar manner. Here, we show that VACV BTB-Kelch protein A55, a close orthologue of EVM150, also binds to Cul3, and this interaction is direct in nature. Surprisingly, the affinity between A55 and Cul3 is much higher than the affinities between human BTB-BACK proteins and Cul3 (Fig.2, Table 1). To understand the molecular basis of this tight interaction the crystal structure of the A55BB/Cul3NΔ22 complex was determined using anisotropic diffraction data extending to 2.3 Å (with an observation:parameter ratio equivalent to that of an isotropic 2.8 Å resolution structure). This is the first reported crystal structure of a virus BTB-Kelch protein in complex with the E3 ubiquitin ligase scaffold protein Cul3.

The overall conformation of the A55/Cul3 complex resembles closely the structures of other cellular BTB/Cul3 protein complexes, with similar mode of dimerization and conserved Cul3-binding interface despite low sequence identities (Fig. 3*C* to *E*,; Fig. S2). The interface area and the number of interface residues at the A55/Cul3 binding interface are comparable to cellular BTB/Cul3 interfaces (Fig. S2, inset table). The conserved φ-X-D/E motif, which was found to be a key contributor to the interaction between SPOP and Cul3, is conserved in A55 (Fig. 4 and Fig. 5*B*) (13). However, mutation of F54 to glutamate at the φ position in A55 only moderately reduced its affinity for Cul3 (Fig. 6, *C* and *D*), whereas the equivalent mutation in SPOP resulted in complete loss of binding (13). A55 residue I48, adjacent to F54, makes more extensive contacts with Cul3 than the equivalent residues in cellular BTB proteins (Fig. 5, *C* to *F*). The I48E mutation completely abolished binding of A55 to the Cul3 N-terminal domain lacking the NTE (Cul3NΔ22) and reduced affinity for the full Cul3 N-terminal domain (Cul3N) by several orders of magnitude (Fig. 6, *G* and *H*; Table 3). An equivalent mutation (A77E) completely abolishes binding of KLHL3 to Cul3NΔ20, further supporting the significance of this hydrophobic interaction for binding of BTB domains to Cul3 (7). Interestingly, ITC studies suggest that the A55/Cul3 interaction is more entropically favourable than the KLHL3/Cul3 interaction (Fig. 2 and Table 1), whereas structural analysis suggests that hydrophobic interactions contribute less energy (via the entropically-favourable release of solvent) to the A55/Cul3 interaction (Fig. S2, inset table). In Cul3-bound KLHL3, KLHL11 and SPOP structures the α3-β4 loops, which contain the φ residue and interact with hydrophobic pockets on the Cul3 surface, adopt helical conformations (Fig. 5, *B*, *D* to *F*). In A55, the φ residue F54 also fits into a deep hydrophobic pocket on the surface of Cul3 but the α3-β4 loop of A55 does not adopt a helical conformation (Fig. 5*C*). Interestingly, in the structures of unbound KLHL11 and SPOP the α3-β4 loop containing the φ residue is less well ordered and has a different conformation (6,8). Such structural rearrangement upon binding to Cul3 would present an entropic penalty to binding. It is tempting to speculate that a lack of such α3-β4 loop rearrangement, rather than the burial of exposed hydrophobic regions, contributes to the entropically-favourable tight binding of A55 to Cul3. Further, such structural rearrangement may not occur in the absence of the favourable hydrophobic interaction mediated by the φ residue, suggesting a mechanism by which binding to Cul3 would be more significantly diminished for SPOP than for A55 when this residue was mutated.

Sequence alignments reveal the BTB-BACK domains of A55 and other orthopoxvirus BTB-Kelch orthologues such as EVM150 to share extensive (>77%) identity (Table 4), including conservation of the residues in the φ-X-D/E motif and I48 (Fig. 7). This strongly suggests that the interaction between A55 orthologues and Cul3 is conserved amongst poxviruses. In contrast, most of the Cul3-binding residues of A55 are not conserved in the other two VACV BTB-Kelch proteins, C2 and F3 (Fig. 7), which share little sequence similarity to A55 (25% and 23% identity, respectively). This suggests that these proteins are unlikely to interact with Cul3, despite being classified as BTB-Kelch proteins, and is consistent with these proteins being functionally distinct (4,31,32). The N-terminal dimerization helix of the BTB domain appears to be missing in C2 (Fig. 7) suggesting that, unlike most BTB-Kelch proteins, C2 may not be able to form homo- or hetero-dimers via the same mechanism as A55.

**Table 4.** Sequence identities between A55BB and the BB domains of poxvirus orthologues and two other VACV BTB-Kelch proteins C2 and F3.

|  | Ectromelia<br>EVM150 | Cowpox<br>A57 | Skunkpox<br>WA-176 | Raccoonpox<br>Herman-172 | Volepox<br>CA-176 | VACV<br>C2 | VACV<br>F3 |
| --- | --- | --- | --- | --- | --- | --- | --- |
| VACV<br>A55 | 95% | 98% | 79% | 77% | 75% | 25% | 23% |

**Figure 7.**
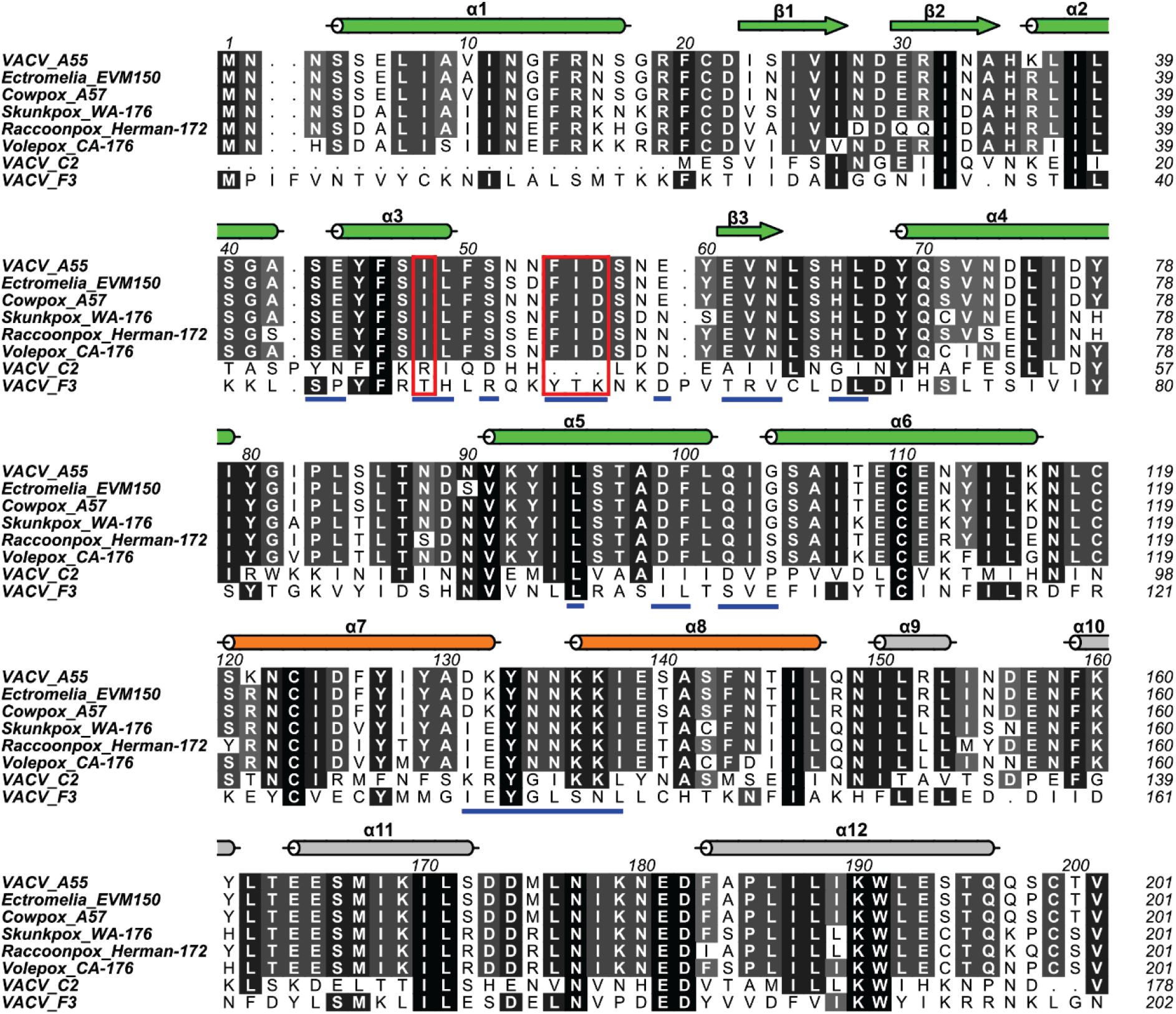
The Cul3-binding residues of A55 are conserved across orthopoxvirus orthologues but not in VACV paralogues C2 and F3. Multiple sequence alignment of the A55 BTB-BACK domains against its paralogues from selected poxviruses and two other VACV BTB-Kelch proteins, C2 and F3. Columns are shaded based on amino acid similarity. Secondary structural elements for A55 are shown above the aligned sequences and colored as in Fig. 3*A*. Residues at the A55/Cul3 interface are underlined in blue. Residues aligned with A55-I48 and the φ-X-E motif are boxed in red.

*In vivo*, VACV expressing A55 induced a smaller lesion in murine model of intradermal infection compared to a virus lacking A55 (4). In cells, the BTB domain of the ECTV EVM150 was reported to inhibit TNFα-induced NF-κB activation; however, this inhibition appears to be Cul3 independent (42). The interaction between A55 and Cul3 therefore is unlikely to be relevant for the inhibition of NF-κB signalling. As a scaffold protein for an E3 ubiquitin ligase complex, Cul3 not only interacts with the BTB-Kelch family of adaptor proteins but also other BTB-domain containing adaptor proteins such as BTB zinc-finger proteins, MATH-BTB proteins, small RhoBTB GTPases and potassium channel tetramerization domain (KCTD) proteins (5,8,10,13,20,24,52). The outcome of the interaction will depend on the specific substrates recruited by the BTB adaptor proteins and will regulate a diverse range of cellular processes, including hypoxic response, ion-channel gating, as well as cytoskeleton organization (19,53,54). The fact that A55 is able to bind Cul3 with much higher affinity than reported cellular binding partners suggest two possible functions of A55. First, A55 may bind to Cul3 and redirect the E3 ubiquitin ligase complex to ubiquitylate otherwise untargeted proteins for proteasomal degradation. Alternatively, A55 may sequester Cul3 and prevent the ubiquitylation and/or proteasomal degradation of proteins that are normally ubiquitylated/degraded upon viral infection. Further experiments are required to discriminate whether A55 fulfils either or both of these roles during infection.

A55 binding to the Cul3 N-terminal domain is significantly increased by the presence of the N-terminal 22 amino acids of Cul3 (Table 1), and the I48E mutant lacks the ability to bind Cu3NΔ22 yet can bind Cul3N with nanomolar affinity (Table 3). This N-terminal extension (NTE) of Cul3 has been shown to interact extensively with a hydrophobic groove formed primarily by the 3-box region of KLHL11 (Fig. S6*A*) (6). Comparison of the structures of KLHL11 in complex with Cul3N or Cul3NΔ22 shows this groove to be pre-formed, rather than being induced by NTE binding (Fig. S6, *A* and *B*). However, sequence and structural alignment of A55 and KLHL11 suggest that an equivalent hydrophobic groove is not present on the surface of A55 (Fig. 4 and Fig. S6*C*). Indeed, the BTB, 3-box and BACK domains are arranged in linear fashion in A55 whereas they form a crescent in KLHL11 or KLHL3 (Fig. S4, *I* to *K*). It is therefore likely that A55 binds the Cul3-NTE via a different set of interactions. Crystallization trials of A55 with Cul3 containing the NTE region (Cul3N) have to date been unsuccessful, and further studies are thus needed to identify residues key for the interaction between A55 and the Cul3-NTE.

## Conclusion

The structure of the first virus BTB-Kelch protein in complex with Cul3 is presented here, which has provided insight into how poxviruses may utilize the host Cul3-based E3 ubiquitin ligase complex for its own benefit. A55 binds Cul3 with much higher affinity than cellular BTB-Kelch proteins. A single point mutation in A55, I48E, significantly diminishes Cul3 binding and could be exploited by future studies to probe the contribution of the A55/Cul3 interaction to VACV virulence.

## Experimental Procedures

### Constructs design

Codon optimized vaccinia virus (VACV) strain Western Reserve (WR) gene *A55R* (Uniprot P24768) full length, *A55R* BTB (residues 1-250) and *A55R* Kelch (residues 251-565), or VACV WR *B14R* (Uniprot P24772) were subcloned into pCDNA4/TO for inducible expression in mammalian cells with an N or C-terminal STREPI and STREPII tag followed by FLAG tag (TAP), respectively. The mammalian expression vectors pcDNA-myc-*CUL3* (19893) and pcDNA-myc-*CUL5* (19895) were purchased from Addgene. The sequence encoding A55 BTB-BACK domain (BB) of the VACV strain WR (residues 1-250) was codon-optimized for expression in mammalian cells and cloned into the pOPTnH vector (55) for expression in *E. coli* with a C-terminal Lys-His_6_ tag. Human Cul3NΔ22 (Uniprot Q13618, residues 23-388) and Cul3N (Uniprot Q13618, residues 1-388) with the I342R and L346D stabilizing mutations (6) in pNIC-CTHF with C-terminal TEV-cleavable His_6_-tags were a gift from Nicola Burgess-Brown (Addgene plasmids 53672 and 53673). KLHL3 (Uniprot Q9UH77, residues 24-276) cloned into pMCSG7 with an N-terminal TEV-cleavable His6-tag was a gift from Alan X. Ji and Gilbert G. Prive (7). Quick-change mutagenesis PCR (Agilent) was used to generate the A55-D56A, A55-F54E, A55-I48E and A55-D99A mutants as per the manufacturer’s protocol.

### Immunoprecipitation

HEK293T-REx (Invitrogen) cells were maintained in Dulbecco’s modified minimal essential medium (DMEM; Gibco) supplemented with 10% foetal bovine serum (FBS; Pan Biotech), non-essential amino acids (NEAA; Gibco) and 50 μg/ml penicillin/streptomycin (Gibco) at 37 °C in a 5% CO_2_ atmosphere. HEK293T-REx inducible cell lines were constructed following transfection with the pCDNA4/TO expression plasmids described above using LT1 transfection reagent following the manufacturer’s instructions (MirusBio). Transfected cells were selected and maintained in DMEM supplemented with 10 μg/ml blasticidin and 100 μg/ml zeocin, following the manufacturer’s instructions (Invitrogen). B14-TAP, TAP-A55, TAP-A55-BTB (TAP-A55BB) or TAP-A55-Kelch HEK293T-REx cells were induced for 24 h with 2 μg/ml doxycycline, washed in ice-cold phosphate-buffered saline (PBS) and subsequently were lysed in either 0.5% NP40 (IGEPAL CA-630) in PBS supplemented with protease inhibitor or RIPA buffer (50 mM Tris pH 8.0, 1% NP40, 150 mM NaCl, 0.5% sodium deoxycholate, 0.5 mM EDTA, 0.1% SDS supplemented with protease inhibitor) where stated. Lysates were cleared at 15,000 *g* at 4 °C and proteins were immunoprecipitated at 4 °C overnight with FLAG M2 beads or Fastflow G-Sepharose (GE Healthcare) incubated previously with mouse monoclonal anti-myc clone 9B11 (CST; #2276) at 1:50 dilution. Beads were washed 3 times in 1 mL lysis buffer by centrifugation for 1 min at 8,000 *g*. After the final wash, beads were incubated in 4× sample loading dye (LDS; Tris 0.5 M pH 6.8, 40% glycerol, 6% SDS, 1% bromophenol blue and 0.8% β-mercaptoethanol), boiled and analysed by immunoblotting.

### Protein expression and purification

Wild-type and mutant A55BB and Cul3NΔ22 were expressed in B834(DE3) *E. coli* cells (Novagen), while Cul3N and KLHL3 were expressed in Rosetta2(DE3)pLysS *E. coli* cells (Novagen). Bacteria were grown at 37 °C in 2× TY medium with shaking at 200 rpm to an OD600 of 0.7-0.9, whereupon protein expression was induced by either adding 0.2 mM IPTG and incubating at 37 °C for 4 h (Cul3N) or by cooling the cultures to 22 °C, adding 0.2 mM IPTG and incubating for 4 h (Cul3NΔ22) or overnight (wild-type and mutant A55). Cells were harvested by centrifugation at 5,000 *g* for 15 min and pellets were stored at -80 °C.

Cells were thawed and resuspended in lysis buffer containing 20 mM HEPES pH 7.5, 500 mM NaCl, 1 mM β-mercatoethanol, 0.05% Tween-20, 0.5 mM MgCl_2_, 400 U bovine DNase I (Roche) and 200 μL EDTA-free protease inhibitor mixture (Sigma-Aldrich). Cells were lysed by passage through a TS series cell disruptor (Constant Systems) at 24 kpsi. Lysates were collected and cleared by centrifugation at 40,000 *g* for 30 min at 4 °C. Cleared lysates were applied to a 5 mL HiTrap TALON crude column (GE Healthcare) pre-equilibrated with binding buffer (20 mM HEPES pH 7.5, 500 mM NaCl, 5 mM β-mercaptoethanol) to capture the His_6_-tagged proteins. The column was washed with binding buffer and the bound proteins were eluted with a gradient 10-150 mM imidazole in binding buffer. Eluted proteins were pooled, concentrated and further purified by SEC using a Superdex 200 column (GE Healthcare) equilibrated in gel filtration buffer (20 mM HEPES pH 7.5, 200 mM NaCl, 1 mM DTT). For Cul3N, an additional anion-exchange chromatography purification step was performed by exchanging the protein into 20 mM Tris pH 7.5, 10 mM NaCl, 1 mM DTT and applying to a MonoQ 5/50 GL column (GE Healthcare) before eluting with a linear gradient of NaCl (10 mM to 1M). Purified proteins were concentrated, snap frozen in liquid nitrogen and stored at - 80 °C. A55BB migrates more rapidly than expected in SDS-PAGE: peptide mass fingerprinting was used to confirm the identity and integrity of the purified protein.

### Size-exclusion chromatography coupled to multi-angle light scattering (SEC-MALS)

SEC-MALS experiments were performed at room temperature. For each experiment, 100 μL protein at 3 mg/mL was injected onto a Superdex 200 increase 10/300 GL column (GE Healthcare) pre-equilibrated with 20 mM HEPES, 150 mM NaCl and 2 mM DTT at a flow rate of 0.5 mL/min. The static light scattering, differential refractive index and the UV absorbance at 280 nm were measured in-line by DAWN 8+ (Wyatt Technology), Optilab T-rEX (Wyatt Technology) and Agilent 1260 UV (Agilent Technologies) detectors. The corresponding molar mass from each elution peak was calculated using ASTRA 6 software (Wyatt Technology).

### Isothermal titration calorimetry (ITC)

ITC experiments were carried out at 25 °C on an automated MicroCal PEAQ-ITC (Malvern Panalytical). Proteins were exchanged into gel filtration buffer (20 mM HEPES, 200 mM NaCl, 1 mM DTT) either by SEC or extensive dialysis prior to experiments. Titrants (wild-type and mutant A55 and KLHL3) at concentrations between 70 and 100 μM were titrated into 7 μM of titrates (Cul3NΔ22 or Cul3N) either as 19 × 2 injections (wild-type A55 and KLHL3) or 13 × 3 injections (mutant A55). Data were analyzed using the MicroCal PEAQ-ITC analysis software (Malvern Panalytical) and fitted using a one-site binding model.

### Reductive methylation

Reductive methylation was carried out at 4 °C using modified protocols from Walter *et al.* (46). Purified A55BB was diluted to 0.8 mg/mL and dialyzed into buffer containing 50 mM HEPES pH 7.5 and 250 mM NaCl. The protein was mixed with 20 μL/mL of 1 M dimethylamine-borane complex (ABC, Sigma) and 40 μL/mL of 1% formaldehyde (UltraPure EM grade, Polysciences) and incubated for 2 hrs at 4 °C. This step was repeated once before mixing with an additional 10 μL/mL of 1 M ABC and incubating overnight at 4 °C. The reaction was quenched with 10 μL of 1 M Tris pH 7.5. Methylated A55BB was further purified by SEC using a Superdex 200 10/300 GL column equilibrated in 20 mM Tris pH 7.5, 200 mM NaCl and 1 mM DTT before being concentrated, snap-frozen and stored at -80 °C.

### Isoelectric focusing (IEF) gel analysis

The IEF gel analysis was performed at 4 °C using a Novex pH 3 to 7 IEF gel (ThermoFisher Scientific) according to manufacturer’s instructions. Native and methylated A55BB were diluted with MilliQ water to 0.8 mg/mL in a total volume of 5 μL and mixed with an equal volume of 2× Novex pH 3 to 10 IEF sample buffer (ThermoFisher) before loading onto the IEF gel. The gel was fixed in 12% TCA for 30 min and washed with MilliQ water before staining with InstantBlue Protein Stain (Expedeon).

### Differential Scanning Fluorimetry (DSF)

DSF experiments were performed in 96-well PCR microplates (Axygen Scientific) on a ViiA 7 Real-Time PCR machine (Life Technologies). To each well of the plate, buffer (20 mM HEPES, 200 mM NaCl, 1 mM DTT), protein and 10× protein thermal shift dye (Applied Biosystems) were mixed at 8:1:1 volume ratio in a final volume of 20 μL and a protein concentration of 0.5 μg/μL. Samples were subjected to thermal denaturation from 25 °C to 95 °C with 1 °C increment per 20 s and real-time fluorescence was recorded. Normalized melt curves were fitted to a biphasic sigmoidal curve using Prism7 (GraphPad Software) and the melting temperatures (Tm) were taken as the mid-point of the first sigmoid.

### Crystallization and data collection

Methylated A55BB was mixed with Cul3NΔ22 at 1:1 molar ratio and the complex was purified by SEC using a Superdex 200 10/300 GL column (GE Healthcare) in 20 mM Tris pH 7.5, 200 mM NaCl, 1 mM DTT. The purified complex was concentrated to 16.3 mg/mL and sitting-drop vapour diffusion experiments were attempted by mixing 100 nL protein with 100 nL reservoir (4% (v/v) tacsimate pH 6.5, 12% (w/v) PEG3350) and equilibrating against 80 μL of the reservoir solution at 20°C. Thin needles were observed after two weeks. Varying the pH, concentration of the tacsimate, concentration of PEG3350 and the protein:reservoir ratio in the sitting drops gave rise to larger crystals that diffracted to ~3.8 Å on Diamond beamline I03. For further optimization, seed stocks for microseeding were generated as described previously (56). Briefly, crystals were crushed and transferred into 50 μL of stabilizing solution (original reservoir solution), vortexed, and seven five-fold serial dilutions of seed into stabilizing solution were generated. Sitting drops were prepared using 100 nL of protein, 150 nL of reservoir and 50 nL of seed stock. Eventually, a drop containing 3.29% (v/v) tacsimate pH 6.5, 9.92% (w/v) PEG3350 and 50 nL of 625-fold diluted seed stock gave rise to crystals that diffracted to 2.3 Å in the best direction on Diamond beamline I04. The crystals were cryoprotected by briefly sweeping through reservoir solution containing 25% (v/v) glycerol and flash-cryocooled by plunging into liquid nitrogen. Diffraction data were collected at 100K on the Diamond beamline I04. Data were indexed and integrated using DIALS (57) as implemented by the xia2 processing pipeline (58). Due to severe anisotropic diffraction, diffraction data were subject to anisotropic scaling using STARANISO (47) and AIMLESS (59).

### Structure determination

The structure of A55BB(M)/Cul3NΔ22 complex was solved by molecular replacement using PHENIX PHASER-MR (60). An initial search using each domain of the SPOP/Cul3 complex (13) (PDB code 4EOZ) as search models successfully placed one copy of Cul3NΔ22, but no solution corresponding to A55BB was forthcoming. MOLREP (61) from the CCP4 program suite (62) was used to locate the A55 BTB domain using B-Cell Lymphoma 6 BTB Domain (49) (PDB code 1R29) as a search model. The 3-box region and the first four helices of the A55 BACK domain (α9-α12) were manually built using COOT (63) with iterative rounds of refinement using Refmac5 (64). The structure was improved by the use of real time molecular dynamics assisted model building and map fitting with the program ISOLDE (50), followed by TLS and positional refinement using BUSTER (51). The quality of the model was monitored throughout the refinement process using Molprobity (65). The atomic coordinates and structure factors (code 6I2M) have been deposited in the Protein Data Bank (http://wwpdb.org/).

### Bioinformatics and structural analysis

Multiple sequence alignments were performed using Clustal Omega (66) and annotated using ALINE (67). Analyses of the binding interfaces were performed using the PDBePISA webserver (68). Molecular figures were generated using PyMOL (69).

## Acknowledgments

We thank Diamond Light Source for access to beamlines I03 and I04 (mx8547), remote access to which was supported in part by the EU FP7 infrastructure grant BIOSTRUCT-X (Contract No. 283570). We thank Janet Deane for assistance with SEC-MALS experiments and useful discussions. We thank Alan X. Ji and Gilbert G. Privé for sharing the pMCSG7-His-TEV-KLHL3 plasmid.

## Footnotes

1 The abbreviations used are: BACK, BTB and C-terminal Kelch; BTB, Bric-a-brac, Tramtrack and Broad-complex; BB, BTB–3-box–BACK domain; Cul3, cullin-3; C3RL, Cul3-RING based E3 ubiquitin ligase complex; DSF, differential scanning fluorimetry; ECTV, ectromelia virus; EV, empty vectors; IEF, isoelectric focusing; ITC, isothermal titration calorimetry; KCTD, potassium channel tetramerization domain protein; K_*D*_, dissociation constant; KLHL, kelch-like protein; MALS, multi-angle light scattering analysis; MATH, Meprin and TRAF homology domain; NTD, N-terminal domain; NTE, N-terminal extension; POZ, poxvirus and zinc finger; RING, really interesting new gene; SEC, size-exclusion chromatography; SPOP, speckle-type POZ protein; TAP, tandem affinity purification tag; TRAF, tumour necrosis factor receptor associated factor; VACV, vaccinia virus.

